# Food web aggregation: effects on key positions

**DOI:** 10.1101/2021.04.18.440319

**Authors:** Emanuele Giacomuzzo, Ferenc Jordán

**Affiliations:** Centre for Ecological Research, Budapest, Hungary; Democracy Institute, Central European University, Budapest, Hungary; Stazione Zoologica Anton Dohrn, Napoli, Italy

**Keywords:** keystone species, centrality indices, trophic role, ecological networks, data aggregation, trophospecies

## Abstract

Providing standard definitions of what should be considered as a node in food webs is still an unsolved problem. Especially for comparative and predictive food web modelling, a more systematic understanding is needed for the effects of trophic aggregation procedures. Aggregation is unavoidable during data management. Therefore, it is crucial to know whether food web properties are conserved during this process.

Here, we study how different aggregation methods change the positional importance of species in food webs. In particular, we investigated the effects of various aggregation algorithms on 24 indices of importance. Our work was carried out on 76 aquatic food webs coming from the Ecopath with Ecosim database (EcoBase). We considered six main types of aggregation, according to the way that the nodes were clustered. These were (i) hierarchical clustering based on the Jaccard index, (ii) hierarchical clustering based on the regular equivalence index (REGE), (iii) maximisation of directed modularity, (iv) maximisation of modularity according to modules in which species fed on the same preys, (v) maximisation of modularity according to modules in which species are fed upon by the same predators, and (vi) clustering through the group model.

Hierarchical clustering based on the Jaccard index and REGE index outperformed the other four methods on maintaining the relative importance of species for all the indices of importance (except for the contrastatus index (*s*′) and betweenness centrality (BC)). The choice between these two methods should follow our research question and the importance index we are interested in studying. The other four aggregation methods change more the centrality of species, especially the one based on maximising directed modularity. When using these aggregation algorithms, one has to keep in mind that the network will not only be smaller but also provides different information.

## Introduction

Trophic data management is something that ecologists always must deal with when working with food webs. Trophic interactions can be described among individuals, life stages, species, higher taxa, functional groups, and several other, appropriately defined nodes of food webs. Some kind of aggregation is unavoidable, and even the most highly resolved food webs contain big aggregates (e.g., “cyanobacteria'', see D’Alelio, Libralato, Wyatt, & Ribera D’Alcalà (2016)). At the same time, even the least resolved food webs may contain species (e.g., “hake”, see Yodzis (1998)). Data aggregation can happen also at later stages, during data analysis, especially in large networks, where the study of hundreds of nodes would be unfeasible (Yodzis & Winemiller, 1999).

Data aggregation methods are problem dependent. Not considering this can bias the way by which we interpret the results of food web models (Hall & Raffaelli, 1993; Paine, 1988). For instance, various levels of aggregation at different trophic levels might bias our interpretation if we are trying to characterise the structure of a network (Yodzis & Winemiller, 1999). Both low- and high-resolution networks can be useful or useless, the key challenge is to properly match the problem, the data management, and the model construction. Even if this seems like a ubiquitous problem in food web ecology, standards for whether and how to aggregate data in a meaningful way does not exist yet.

The process of data aggregation assumes that there are nodes in the network that are similar enough that we can consider them functionally equivalent. For example, two fishes from the same genus might be aggregated into a node of the genus (e.g., *Poecilia sphenops* and *Poecilia reticulata* could be aggregated into *Poecilia*). Similarity can be understood mathematically (equivalent network positions) and biologically (similar trophic habits). Yodzis & Winemiller (1999) and Luczkovich et al. (2003) tried to find nodes in equivalent network positions by borrowing two definitions from social networks. Yodzis & Winemiller (1999) borrowed the concept of structural equivalence – where two nodes are similar when sharing a high number of neighbours – and called the aggregation of structurally equivalent species” tropho-species”. Luczkovich et al. (2003) borrowed the concept of regular equivalence – where two nodes are similar when sharing a high number of similar but not necessarily the same neighbours. Nodes belonging to the same equivalence class are said to be sharing the same trophic role.

Groups of nodes that have different neighbours but form dense subgraphs are called modules. Species in food web modules can play different roles (e.g., predator and prey), but they maintain well-defined multispecies processes (e.g., connecting benthic and pelagic organisms). Aggregating the modules of a food web has been suggested already by Allesina & Pascual (2009). The two most reliable ways of finding modules in food webs are through the group model and modularity maximisation. The group model was firstly developed by Allesina & Pascual (2009) and then made more computationally efficient by Sander, Wootton, & Allesina (2015). Modularity maximisation was applied to food webs by Guimerà et al. (2010) following three definitions of modularity. The first one, which we will refer to as density-based modularity, is the degree to which nodes inside modules interact more among themselves than with nodes of other modules. The second one, which we call prey-based modularity, is the degree to which nodes inside modules tend to interact with the same predators. The third one, which we gave the name of predator-based modularity, is the degree to which nodes inside modules tend to interact with the same preys.

The positional importance of species differs in both highly aggregated and highly resolved networks. Central positions may be a proxy for functional importance and the community-wide distribution of either centrality values (Bauer, Jordán, & Podani, 2010) or hypothetical importance values (Mills, Doak, & Soulé, 1993) provide macroscopic descriptors of ecosystems. Here, we investigate how different aggregation methods maintain the relative importance of species according to 24 of the most used indices of importance in food web research. Our investigation was carried out on 76 Ecopath with Ecosim (EwE) food web models available on the EcoBase database (Colléter et al., 2013). By having been constructed with the same methodology (see Okey (2004)), they provided us with comparable results. See supporting information for a list of these food webs.

## Material and methods

### Clustering techniques

To cluster similar nodes, we used the following six clustering techniques.

#### Hierarchical clustering with Jaccard index

As a first clustering method, we clustered structurally equivalent nodes as in Yodzis & Winemiller (1999). We used the Jaccard similarity index (Jaccard, 1912) as a measure of structural equivalence. See supporting information for the clustering algorithm.

#### Hierarchical clustering with REGE index

Our second clustering method consisted of clustering regularly equivalent nodes as in Luczkovich et al., (2003). The measure of regular equivalence we used was the REGE index (Borgatti & Everett, 1993). See supporting information for the clustering algorithm.

#### Clustering of density-based modules

As a third clustering method, we clustered the nodes inside the modules found by maximising the density modularity, as in Guimerà et al. (2010). This type of modularity is expressed as the number of extra links present within the modules compared to the ones expected by chance. For directed networks, it can be expressed through the following equation of Arenas, Duch, Fernández, & Gómez, (2007), which is a generalisation of the Newman-Girvan modularity (Newman, 2004)

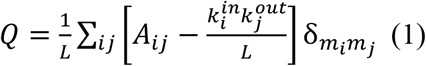

where *Q* is the modularity of the network, *L* is the number of links in the network, *A_ij_* is the element of the adjacency matrix of the directed binary network (links go from *j* to *i*), 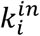 is the indegree of *i*, 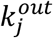 is the outdegree of *j*, *m_i_* is the module of *i*, *m_j_* is the module of *j* and δ is the Kronecker delta (Kozen & Timme, 2007).

The number and composition of the modules were found by using the Leiden algorithm of Traag, Waltman, & van Eck (2019). This algorithm is an extension of the Louvain algorithm (Blondel, Guillaume, Lambiotte, & Lefebvre, 2008). The latter is one of the best performing and fastest for community detection (Traag et al., 2019). However, it tends to produce communities that are arbitrarily poorly connected to each other and sometimes even disconnected. The Leiden algorithm not only solves this problem by producing better connected communities, but it is also faster. The code that we used was implemented in the igraph package (Csardi & Nepusz, 2006) for the statistical software R (R Development Core Team, 2013).

#### Clustering of prey-based and predator-based modules

As the fourth and fifth clustering methods, we clustered the nodes of every module that was found by maximising the prey modularity and the predator modularity of the food web, as in Guimerà et al. (2010). In this case, the modularity of the food web is expressed as to how much different nodes connect to the same predators (for prey modularity) or preys (for predator modularity) than expected by chance. Mathematically, it can be expressed by the following equation (Guimerà, Sales-Pardo, & Amaral, 2007) for prey modularity

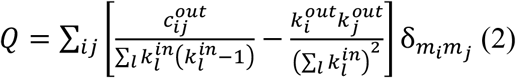

or in the following one for predator modularity

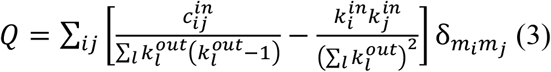

where 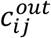 is the number of outgoing links that *i* and *j* have in common and 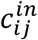 is the number of incoming links that *i* and *j* have in common. We maximised these two types of modularity by using the rnetcarto package (Doulcier & Stouffer, 2015) implemented in R. This finds the community structure of the network by using the optimisation method of simulated annealing (Kirkpatrick, Gelatt, & Vecchi, 1983).

#### Clustering of groups

As a sixth clustering method, we clustered the nodes inside the modules found by the group model of Allesina & Pascual (2009). This model finds the modules that maximise the probability of randomly retrieving the food web by generating a modular version of an Erdős-Rényi random graph. For an arbitrary number of groups *k*, the probability of retrieving the food web is:

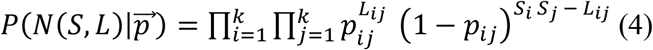

where *N*(*S*, *L*) is the food web *N* with *S* number of nodes and *L* number of links, 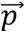 is the vector containing the probabilities of a connection between and within clusters, *p_ij_* is the probability that a node inside the group *i* connects to another node inside the group *j*, *L_ij_* is the number of links connecting nodes belonging to the group *i* to nodes belonging to the group *j*, *S_i_* is the number of nodes in the cluster *i*, and *S_j_* is the number of nodes in the cluster *j*.

Because of the high number of possible module arrangements, it is not possible to explore them all. To find the best possible solution that our computation power allows us to find, we used the algorithm of Sander, Wootton, & Allesina, 2015. This relies on a Metropolis-Coupled Markov Chain Monte Carlo (*MC*^3^), also known as parallel tempering (Geyer, 1991), with a Gibbs sampler (Yildirim, 2012). *MC*^3^ can be considered as a Markov chain Monte Carlo (MCMC) with multiple chains running all at once (Sander et al., 2015).

### Connecting the clusters and assigning interaction strength

The connection of the clusters followed a similar approach to the one described in Martinez (1991). We used five methods to decide whether there was a link between two clusters. The first one produces the maximum connectance and is known as maximum linkage method (NMAX). Here, a cluster has a connection to another cluster if it has at least one link going from one of its nodes to the nodes of the second cluster. The second one produces the minimum connectance and is known as minimum linkage method (NMIN). In this case, a cluster is connected to another only if all its nodes have a connection to all the nodes of the other cluster. The other three methods produce an intermediate connectance. They consider a link from a cluster to another only if at least 25%, 50%, or 75% of possible connections from the first cluster to the second are realised.

The weight of the link was then calculated in four different ways: as the minimum weight, the maximum weight, the mean weight, and the sum of the weights of the links going from the members of the first cluster to the ones of the second cluster.

### Indices of importance

For each food web, we calculated the indices of importance before and after the aggregation. The indices of importance of a node after the aggregation was defined as the one of its cluster. Let’s consider the following example. Before the aggregation, the node “hake” has a degree centrality of 5. The aggregation process clusters it with other fish nodes, creating a node in the aggregated food web called “fish”. The degree centrality of “fish” is 8. In this case, the degree centrality of “hake” is 5 in the original food web and 8 in the aggregated food web. The importance indices we used belonged to the following families.

#### Degree centrality (DC)

The degree centrality (*DC*) of a node *i* is the number of links it has (Wasserman & Faust, 1994)

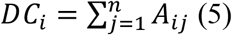

where *n* is the number of nodes in the food web, and *A*_*ij*_ is the element of the adjacency matrix, after the network has been transformed in a binary undirected one.

Another type of degree centrality that we considered was the weighted degree centrality (*wDC*), often referred to as node strength. Its formula is the same as for the non-weighted degree centrality. This time, however, the adjacency matrix is of an undirected weighted network (Fornito, Zalesky, & Bullmore, 2016)

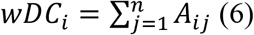

#### Closeness centrality (CC)

The closeness centrality (*CC*) of a node is the average distance of a node from all the others in the network (Wasserman & Faust, 1994)

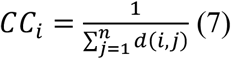

where *d*(*i*, *j*) is the shortest path between node *i* and *j*.

#### Betweenness centrality (BC)

The betweenness centrality (*BC*) of a node is the average number of times that it acts as a bridge along the shortest path between two other nodes. It can be mathematically expressed as follows (Wasserman & Faust, 1994)

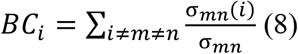

where σ_*mn*_ is the total number of shortest paths going from *m* to *n* and σ_*mn*_(*i*) is the total number of these paths passing through *i*.

#### Status index (s)

The status index of a node is the sum of its distances from all the other nodes inside the network, calculated as their shortest paths following a bottom-up direction (Endrédi, Senánszky, Libralato, & Jordán, 2018)

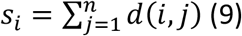

where *d*(*i*, *j*) is the shortest path between node *i* and *j*. It was first introduced to social networks, followed two years later by its application to food webs by Harary (1959, 1961). By following the same method but in a top-down direction we obtain the contrastatus 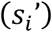

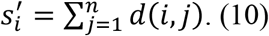

The difference between the status and the contrastatus is called the net status (Δ*s_i_*)

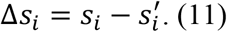

The computation of the status, contrastatus and net status needs to be performed on a network without cycles. See supporting information for the algorithm used to create a directed acyclic graph (DAG).

#### Keystone index (K)

The keystone index was firstly introduced by Jordán, Takacs-Santa, & Molnar (1999) and inspired by the status index. As in the status index family, the keystone index of a species is calculated by considering the bottom-up and the top-down effects separately (Jordán, Liu, & Davis, 2006). Unlike the status index, which only considers the distance between a node and all the other nodes, the keystone index takes into consideration how the size of a certain effect gets split between the different neighbours of a node. Every time the effect reaches a certain node connected to multiple nodes, the following nodes receive only a fraction of the total effect. For example, when considering the bottom-up effect, if the prey has two predators, the bottom-up effect received by each predator will be half. The computation of the keystone index, as the status index, also needs to be performed on a network without cycles.

The keystone index of a species *i* is equal to the sum of its bottom-up and top-down effects

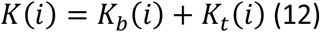

where *K*(*i*) is the keystone index, *k_b_*(*i*) is the bottom-up keystone index, and *k_t_*(*i*) is the top-down keystone index. The bottom-up keystone index is calculated as

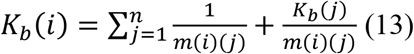

where *j* is a predator of *i* and *m*(*i*)(*j*) is the number of preys of. 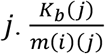 is the fraction of bottom-up effects of *j* that are caused by *i*. The *k_b_*(*j*) of top predators is set as 0. The top-down keystone index is calculated exactly as the bottom-up keystone index, but with the direction of the links inverted.

When calculating the keystone index, we might be also interested in how much of the effect of a species on the community depends upon its directed and undirected effects. To do this, we can split the keystone index of a species into its direct and indirect effect components

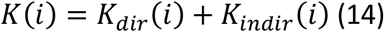

where *K_dir_*(*i*) is its directed component, and *K_indir_*(*i*) is its indirected component. To calculate them, we need to know that

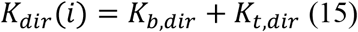

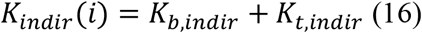

where K_*b,dir*_ is the directed component of the bottom-up effect, K_*t,dir*_ is the directed component of the top-down effect, K_*b,indir*_ is the indirected component of the bottom-up effect, and K_*t,indir*_ is the indirected component of the top-down effect. The directed and indirected components of the bottom-up index are calculated as

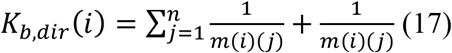

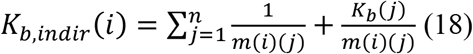

The direct and indirect components of the top-down effect are calculated in the same way, but with the direction of the links inverted.

#### Topological importance (TI)

The topological importance (*TI*) of a node represents its potential to create bottom-up effects on other species, up to a certain number of steps that we can set. It was first introduced to host-parasitoid networks by Müller, Adriaanse, Belshaw, & Godfray (1999) and then to food webs by Jordán, Liu, & van Veen (2003). If topological importance takes interaction strength into consideration, we refer to it as weighted topological importance (*WI*). See supporting information for the algorithm for the computation of topological importance and weighted topological importance.

#### Species uniqueness (STO)

Species uniqueness (*STO*) represents how redundant the strong interactions of a node are. It was first introduced by Lai, Liu, & Jordán (2015) and can be considered as an extension of the trophic field overlap (*TO*) (see Jordán, Liu, & Mike (2009). The trophic field overlap of a node *i* is the number of times that *i* and another node interact strongly with the same predator. Interactions are considered strong when they exceed a certain threshold. To avoid having to choose a single threshold, *TO* is calculated based on multiple thresholds. Each of these *TO* is then summed to give species uniqueness. If we calculate the topological overlap taking into consideration interaction strength, we can calculate weighted species uniqueness (*wSTO*). See supporting information for the algorithm for the computation of trophic field overlap.

## Statistical analysis

The combination of the six clustering techniques, five linkage methods and four ways of determining interaction strength produced 120 ways of aggregating food webs. For each of these aggregation methods, we studied their effects on the indices of importance. In particular, we studied the correlation between the ranking of the nodes before and after the aggregation. This correlation was calculated by using Kendall’s tau b (τ_*B*_) - a version of Kendall’s rank correlation coefficient that makes adjustments for ties (Agresti, 2012). For each combination of aggregation method and index of importance, we found the mean τ_*B*_ across all food webs. This required us to convert τ_*B*_ using the Fisher z-transformation (Fisher, 1915). For each fisher’s z mean, we found its 95% confidence interval by bootstrapping (DiCiccio & Efron, 1996). The fisher’s z means, and 95% confidence intervals were then back transformed to τ_*B*_. τ_*B*_ and bootstrapping were implemented in the Statistics and Machine Learning Toolbox for MATLAB (Mathworks Inc., 2019).

## Results

### Size of the clusters produced

The 76 food webs we used had a median of 25.5 nodes (IQR = 16.0), with a minimum of 14 nodes and a maximum of 55 nodes (Figure 1). The median size of the aggregated food web compared to the original one was 74.5% (IQR=10.8%) for the Jaccard index, 73% (IQR=7.2%) for the REGE index, 12.8% (IQR=6.5) for the density-based modules, 35.8% (IQR=21.3%) for the prey-based modules, 72.1% (IQR=29.6%) for the predator-based modules and 15.8% (IQR=6.5%) for the group model (Figure 2 and 6).

**Figure 1.**
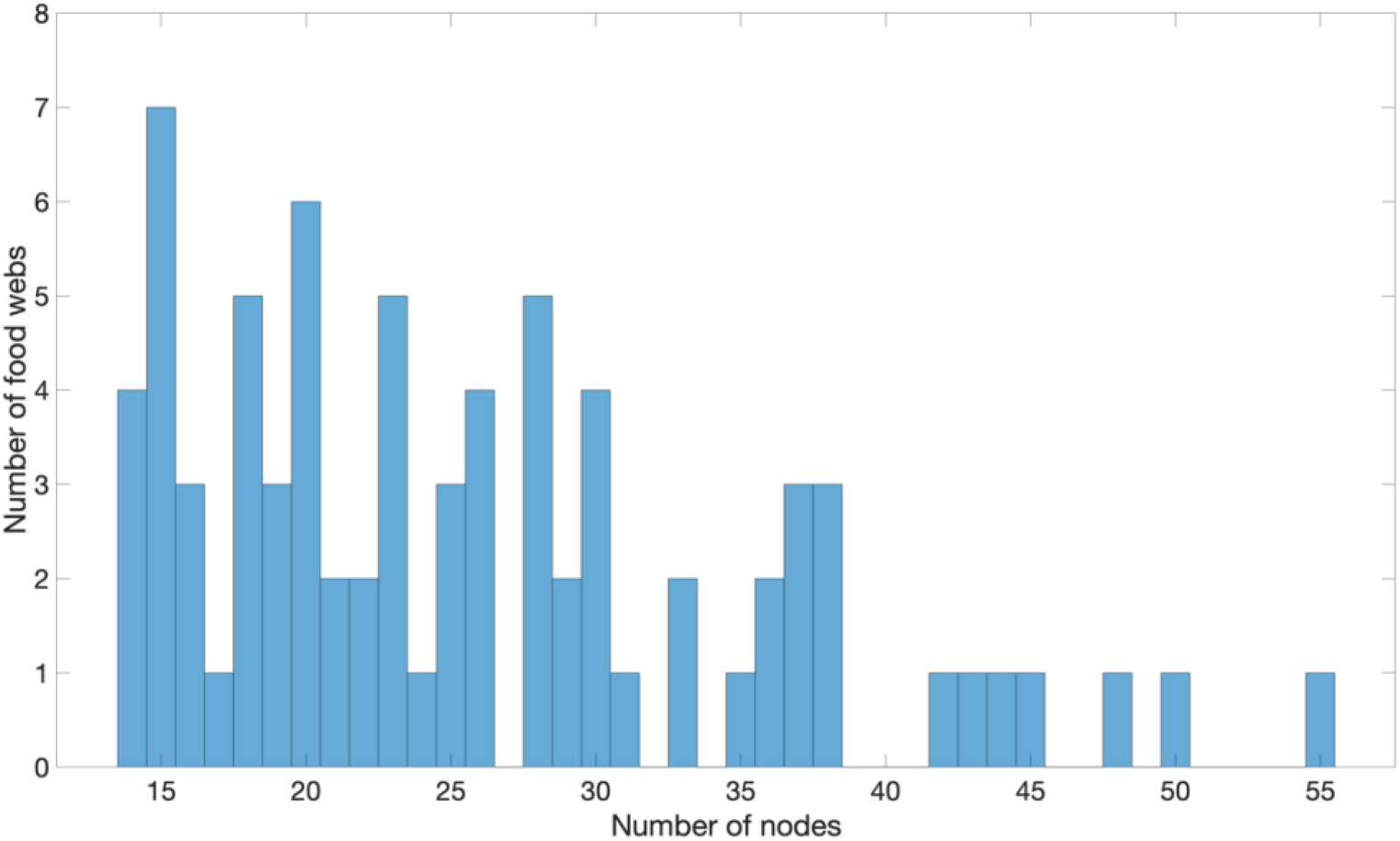
Size of the food webs we used in our study.

**Figure 2.**
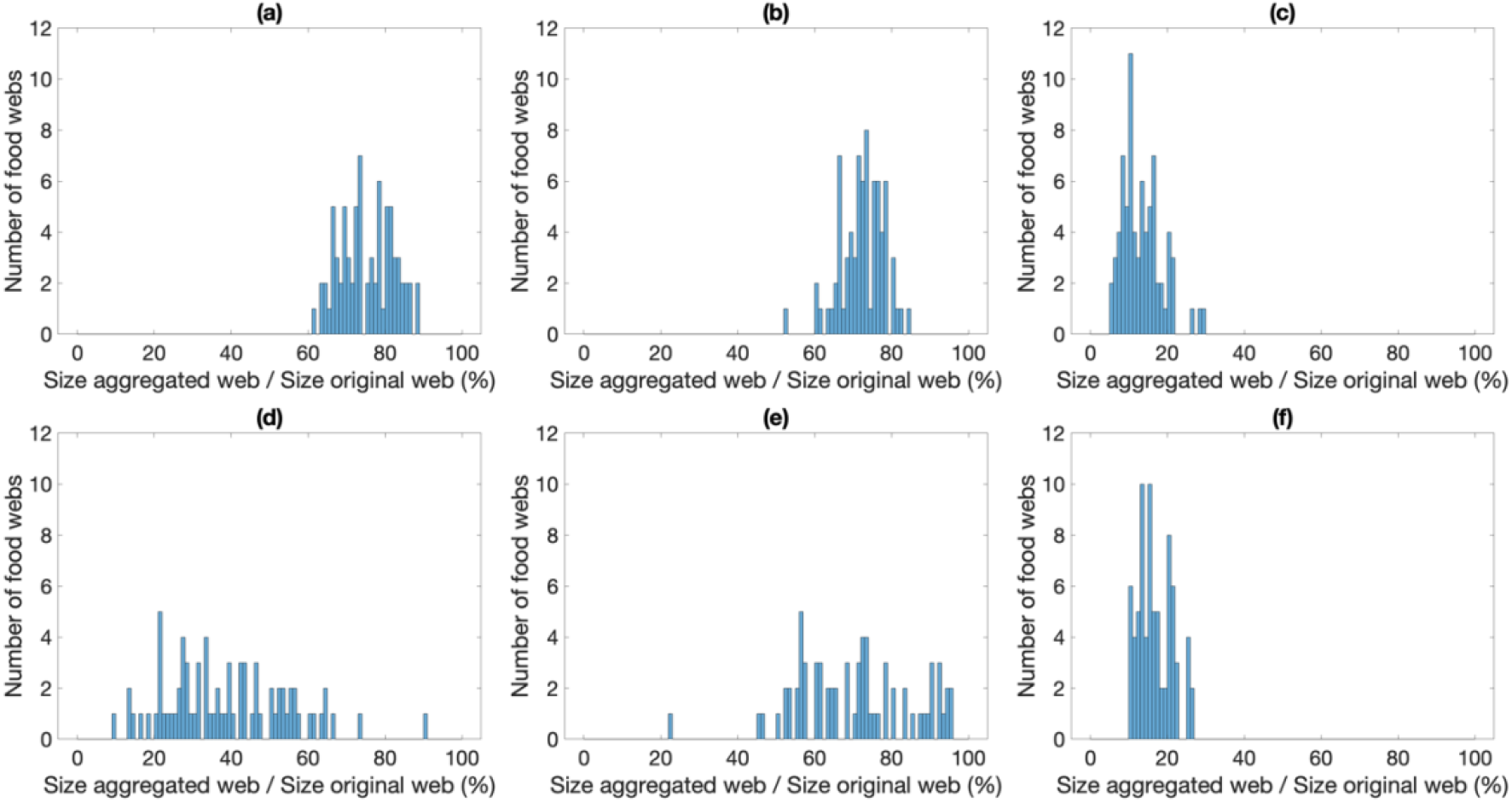
Size of the clusters produced by the different clustering methods. (a) = hierarchical clustering with Jaccard index, (b) = hierarchical clustering with REGE index, (c) = maximisation of density-based modularity, (d) = maximisation of prey-based modularity, (e) = maximisation of predator-based modularity, (f) = group model.

### Correlation of indices of importance before and after the aggregation

The correlation between the ranking before and after the aggregation can be seen in Figure 3. By focusing only on the clustering method and ignoring the linkage method and the interaction strength method, we can select the best clustering for each combination of centrality indices and clustering methods. This provides us with a clearer heat map. See Figure 4. Ranking the clustering algorithms in Figure 4 produces Table 1. Density modularity always ranked as the worst clustering algorithm. Prey-based modules and group model were always ranked as either fourth or fifth. Except for *BC* and *s′*, the clustering of predator modules ranked consistently as third. Excluding the results of contrastatus, the hierarchical clustering based on Jaccard index and REGE index were always ranked as the best clustering methods. Jaccard index was better than REGE for weighted and unweighted species uniqueness, unweighted topological importance, degree centrality, closeness centrality, and betweenness centrality. REGE was better for weighted topological importance and weighted degree centrality. Status index and keystone index were maintained better either by Jaccard or by REGE according to which index of those two families we considered. We can qualitatively say that the correlation between the ranking before and after the aggregation seems to increase with the size of the aggregated food web.

**Table 1.** Best Kendall’s correlation coefficients, as in Figure 4. This time, they are ranked from the clustering that produced the best correlation, to the clustering that produced the worst correlation. Green = hierarchical clustering with Jaccard index, red = hierarchical clustering with REGE index, grey = density-based modules, blue = predator-based modules, purple = groups produced by the group model. C1 = Best clustering, C2 = second best clustering, C3 = third best clustering, C4 = fourth best clustering, C5 = fifth best clustering, C6 = sixth best clustering.

| Index | C1 | C2 | C3 | C4 | C5 | C6 |
| --- | --- | --- | --- | --- | --- | --- |
| *DC* | 0.76 | 0.73 | 0.68 | 0.42 | 0.40 | 0.11 |
| *wDC* | 0.93 | 0.85 | 0.80 | 0.63 | 0.57 | 0.27 |
| *CC* | 0.75 | 0.72 | 0.67 | 0.40 | 0.40 | 0.06 |
| *BC* | 0.77 | 0.72 | 0.71 | 0.46 | 0.33 | 0.00 |
| *s* | 0.94 | 0.87 | 0.85 | 0.79 | 0.74 | 0.23 |
| *s'* | 0.89 | 0.89 | 0.87 | 0.77 | 0.71 | 0.27 |
| $\Delta s$ | 0.90 | 0.88 | 0.86 | 0.77 | 0.74 | 0.25 |
| *k* | 0.72 | 0.72 | 0.62 | 0.32 | 0.28 | 0.04 |
| *k<sub>bu</sub>* | 0.91 | 0.86 | 0.79 | 0.73 | 0.72 | 0.19 |
| *k<sub>td</sub>* | 0.82 | 0.81 | 0.78 | 0.69 | 0.53 | 0.24 |
| *k<sub>dir</sub>* | 0.68 | 0.66 | 0.58 | 0.25 | 0.21 | 0.05 |
| *k<sub>indir</sub>* | 0.74 | 0.73 | 0.65 | 0.43 | 0.40 | 0.04 |
| *TI*<sup>1</sup> | 0.82 | 0.79 | 0.68 | 0.49 | 0.47 | 0.15 |
| *TI*<sup>3</sup> | 0.87 | 0.81 | 0.73 | 0.56 | 0.54 | 0.18 |
| *TI*<sup>5</sup> | 0.88 | 0.82 | 0.74 | 0.58 | 0.55 | 0.20 |
| *WI*<sup>1</sup> | 0.82 | 0.79 | 0.68 | 0.49 | 0.47 | 0.15 |
| *WI*<sup>3</sup> | 0.87 | 0.81 | 0.73 | 0.56 | 0.54 | 0.18 |
| *WI*<sup>5</sup> | 0.88 | 0.82 | 0.74 | 0.58 | 0.55 | 0.20 |
| *STO*<sup>1</sup> | 0.90 | 0.79 | 0.74 | 0.61 | 0.59 | 0.07 |
| *STO*<sup>3</sup> | 0.90 | 0.78 | 0.72 | 0.59 | 0.59 | 0.06 |
| *STO*<sup>5</sup> | 0.88 | 0.77 | 0.72 | 0.59 | 0.58 | 0.07 |
| *wSTO*<sup>1</sup> | 0.87 | 0.81 | 0.72 | 0.59 | 0.57 | 0.10 |
| *wSTO*<sup>3</sup> | 0.85 | 0.82 | 0.71 | 0.59 | 0.53 | 0.11 |
| *wSTO*<sup>5</sup> | 0.84 | 0.83 | 0.70 | 0.59 | 0.51 | 0.11 |

**Figure 3.**
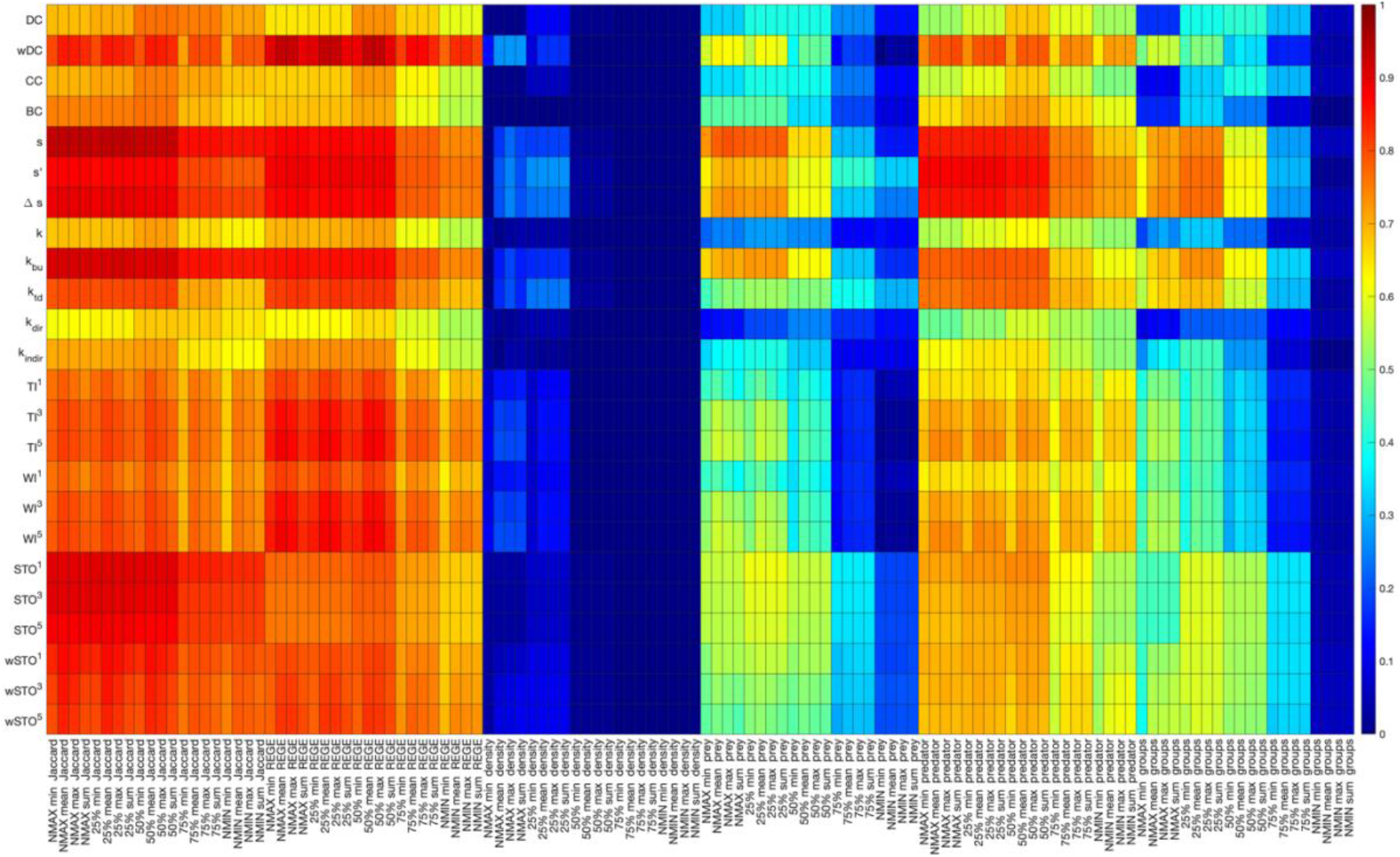
Kendall’s rank correlation (τ) between the ranking of the nodes in the original food web and in the aggregated food web. The x-axis describes the aggregation method, which is composed of three components: (i) clustering algorithm; (ii) linkage method; and (iii) and interaction strength method. Jaccard = hierarchical clustering using Jaccard index, REGE = Hierarchical clustering using REGE index, density = clustering of density-based modules, prey = clustering of prey-based modules, predator = clustering of predator-based modules, groups = clustering of groups. NMAX = maximum linkage method, 25% = at least 25% of links realised to consider a connection, 50% = at least 25% of links realised to consider a connection, 75% = at least 75% of links realised to consider a connection, NMIN = minimum linkage method. min = minimum interaction strength, max = maximum interaction strength, sum = sum of interaction strengths, mean = mean interaction strength. The y-axis describes the importance indices. *DC* = degree centrality, *wDC* = weighted degree centrality, *CC* = closeness centrality, *BC* = betweenness centrality, *s* = status index, *s′* = contrastatus index, Δ*s* = net status index, *k* = keystone index, *k_bu_* = bottom-up keystone index, *k_td_* = top-down keystone index, *k_dir_* = directed keystone index, *k_indir_* = indirected keystone index, *TI*^1^ = 1-step topological importance, *TI*^3^ = 3-step topological importance, *TI*^5^ = 5-step topological importance, *WI*^1^ = 1-step weighted topological importance, *WI*^3^ = 3-step weighted topological importance, *WI*^5^ = 5-step weighted topological importance, *STO*^1^ = 1-step species uniqueness, *STO*^3^ = 3-step species uniqueness, *STO*^5^ = 5-step species uniqueness, *wSTO*^1^ = 1-step weighted species uniqueness, *wSTO*^3^ = 3-step weighted species uniqueness, *wSTO*^5^ = 5-step weighted species uniqueness.

**Figure 4.**
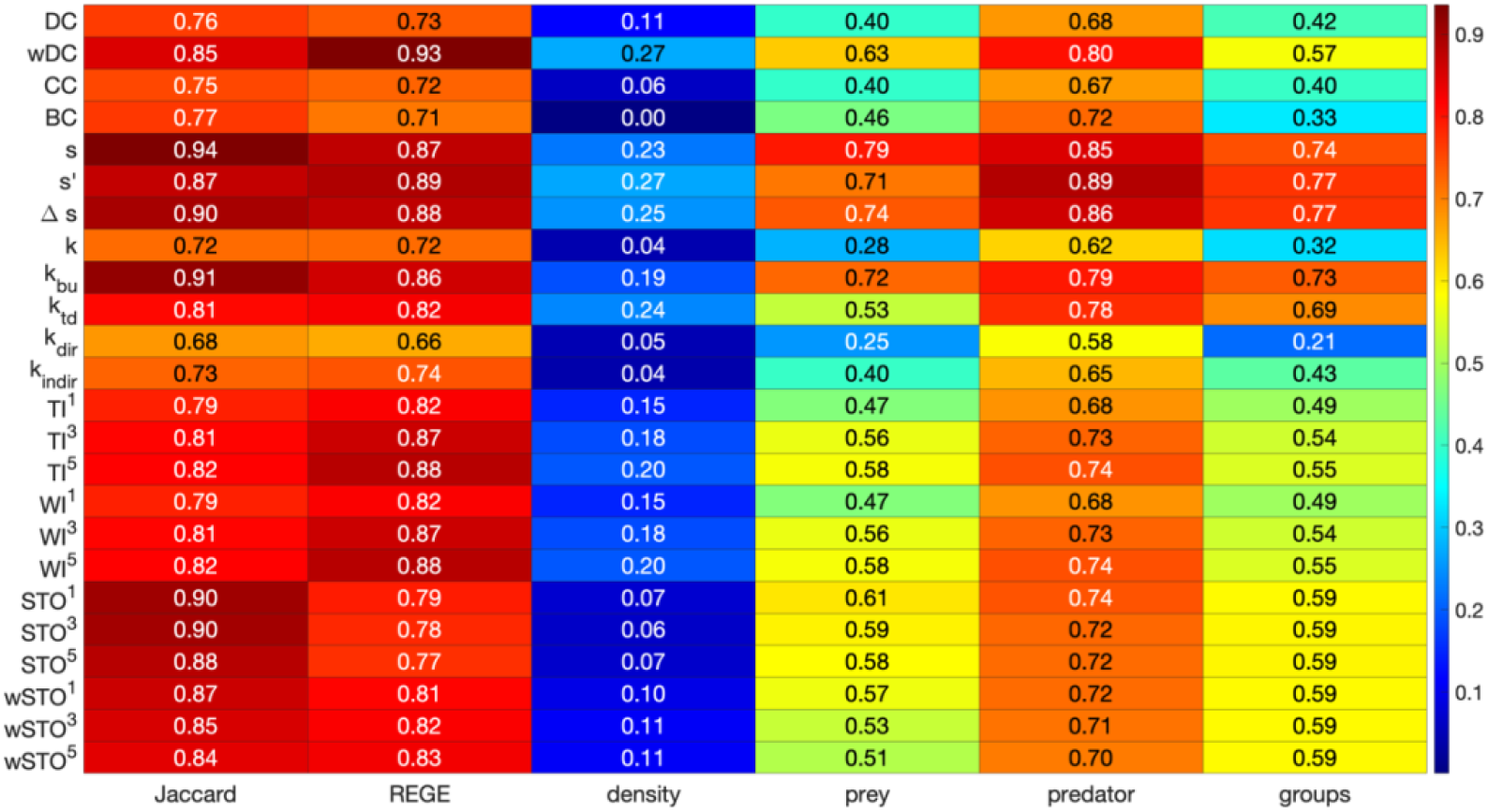
Heat map of the best Kendall’s rank correlation coefficient for each combination of clustering methods and centrality indices. The best correlation is selected across linkage methods and methods of determining interaction strength. Jaccard = hierarchical clustering using Jaccard index, REGE = Hierarchical clustering using REGE index, density = clustering of density-based modules, prey = clustering of prey-based modules, predator = clustering of predator-based modules, groups = clustering of groups. For the confidence intervals associated with these values and for the linkage method & interaction strength method used, see supporting information.

## Discussion

The aggregation of nodes in food webs is an old problem, dating back at least to the beginning of the 1980s (e.g., Pimm (1982)). Multiple studies investigated the effect of different types of aggregation on food web properties, such as connectance, food chain length and ratio between bottom, intermediate and top species (Hall & Raffaelli, 1991; Martinez, 1991, 1993; Sugihara, Bersier, & Schoenly, 1997; Sugihara, Schoenly, & Trombla, 1989). Not much attention, however, has been given to the effects that aggregation has on our ability to find keystone species inside the food web. Essington & Plagányi, (2014) and Plagányi & Essington, (2014) tried to investigate this topic. The former by studying changes in degree centrality and in the ratio between the consumer species biomass and the total consumer biomass. The latter by the SURF index (an index used to find important forage fish species). These, however, focused on indices that are rarely used in keystone species research and did not compare the effect of different aggregation methods.

In this paper, we study the effects of aggregation on several indices used in network ecology (Estrada, 2007; Olmo Gilabert et al., 2019). The aggregation methods that we compared were the widely used hierarchical clustering using Jaccard index and REGE index, as well as directed modularity maximisation, prey-based modularity maximisation, predator-based modularity maximisation, and clustering through the group model. The latter four are normally used for community detection, but we explored the possibility of using them in the future for data aggregation.

Our results show that different aggregation methods maintain the relative importance of species in different ways. Therefore, they have different potential of changing the keystone species of the food web (Figure 5). Except for the contrastatus index (*s′*), hierarchical clustering with the Jaccard index and hierarchical clustering with the REGE index outperformed the other methods. When choosing between these two methods, however, we need to consider that not all indices have the same power to predict keystone species. Gouveia, Móréh, & Jordán (2021) looked at topological indices and how their findings correlated with the findings of the dynamical index keystoneness (Libralato, Christensen, & Pauly, 2006). They found that the most reliable topological index was the weighted degree centrality (*wDC*). This could predict the most important species for dynamic processes in 70.1% of the cases. It was followed by a combination of *wDC* and the 5-step weighted topological importance (*WI*^5^), which increased this percentage to 78.4%. In light of these findings, REGE might be considered the best clustering algorithm, as it maintains *wDC* and *WI*^5^ the best.

**Figure 5.**
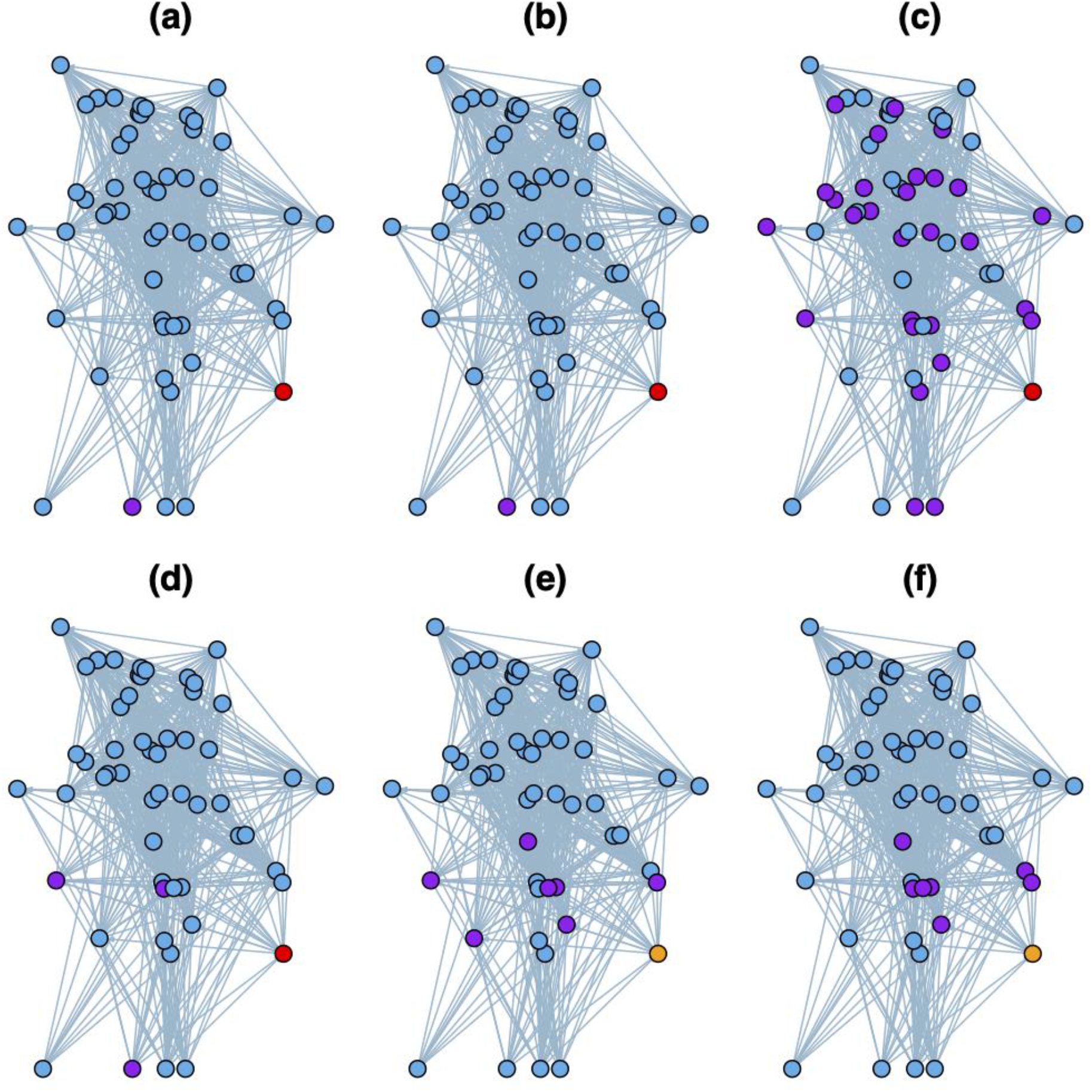
Effect of aggregation on most the most important nodes in a food web. Red: central before and central after. Orange: central before and not central after. Purple: not central before and central after. Blue: not central before and not central after. The self-loops are not included in the figure for clarity. (a) = hierarchical clustering using Jaccard index, (b) = hierarchical clustering using REGE index, (c) = clustering of density-based modules, (d) = clustering of prey-based modules, (e) = clustering of predator-based modules, (f) = clustering of groups. The food web here depicted is the one of the West Florida Shelf (Thomas A. Okey, 2004). It is largest network used in this study (55 nodes). To make the figure clearer, we did not include the self-loops.

The choice of the aggregation algorithm, however, boils down to our research question. The particular index we are interested in might drive the choice between the Jaccard index and the REGE index. Another factor influencing our choice is the resulting network size. How much do we need to aggregate? In case we might want to have higher aggregation, we might consider using the group model, which produces high aggregation, but performs way better than density-based modularity (Figure 6). But also, we might be interest in a particular species role – in that case we should aggregate according to that specific role. Each method reveals some kind of biological similarities between nodes. The key point is to know the effects of these aggregation procedures and to keep them in mind when applying them and evaluating the properties of the aggregated network.

**Figure 6.**
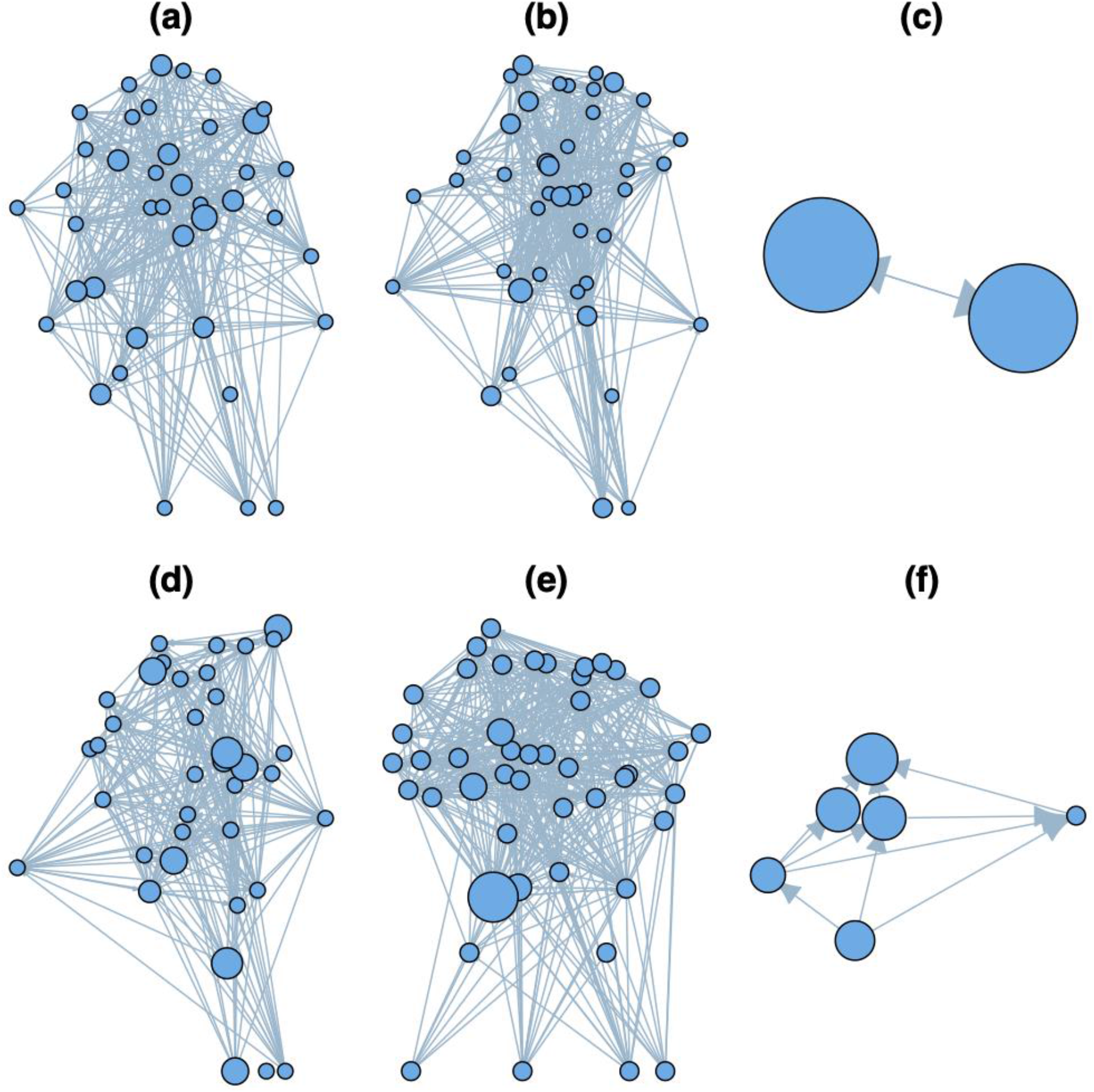
Aggregation of the food web of Figure 5 according to different clustering algorithms. (a) = hierarchical clustering using Jaccard index, (b) = hierarchical clustering using REGE index, (c) = clustering of density-based modules, (d) = clustering of prey-based modules, (e) = clustering of predator-based modules, (f) = clustering of groups. The linkage method and the interaction strength method used for each clustering were the ones that produced the highest Kendall’s rank correlation between the ranking before and after the aggregation. The size of the nodes is proportional to the number of nodes that have been aggregated into them. Also in this case, we did not include self-loops.

In this paper, we focus on a well-defined set of food webs that are methodologically strictly comparable (all created by the EwE methodology). Studying aggregation would be more exciting for larger food webs but these are rarely comparable (lacking standards for their description). When data will be available, interaction networks representing various interaction types will be very important to analyse from our present perspective: different interaction types may be differently sensitive to aggregation. A major advancement would be the analysis of aggregation effects on dynamical food web models. To see how dynamical properties can be altered or changed, as an effect of aggregation algorithms, would be a major step towards predictive food web modelling.

In conclusion, we have shown that different aggregation methods maintain differently the relative importance of species in a food web. Hierarchical clustering with Jaccard index and hierarchical clustering with REGE index have been shown to be the best at doing this. The choice between these two algorithms should depend upon the type of importance index we are interested in maintaining. Future research should be carried out on larger food webs, dynamical indices and determining what the best linkage method and new interaction strength method are.

## Supporting information

Description of Algorithms

## Acknowledgements

We would like to thank Wei-Chung Liu for providing the code for computing some centrality indices, Stefano Allesina & Elizabeth Sander for providing the code for the computation of the group model and Anett Endrédi for providing us support with data management. Ferenc Jordàn was supported by AtlantECO (H2020 BG-08-2018-2019, grant no. SEP-210591007).

## Supplementary material

The code used for this analysis is available at https://github.com/EmanueleGiacomuzzo/Data_aggregation. This work resulted also in the MATLAB toolbox “Food Web Analysis Toolbox” available at https://uk.mathworks.com/matlabcentral/fileexchange/89907-food-web-analysis-toolbox

