## Supplemental Tables for "Food web aggregation: effects on key positions"

| Index | C1 | C2 | C3 | C4 | C5 | C6 |
|---|---|---|---|---|---|---|
| $DC$ | 0.74-0.78 | 0.70-0.75 | 0.63-0.72 | 0.36-0.46 | 0.36-0.47 | 0.09-0.15 |
| $wDC$ | 0.93-0.94 | 0.84-0.86 | 0.76-0.83 | 0.59-0.66 | 0.54-0.61 | 0.24-0.32 |
| $CC$ | 0.73-0.77 | 0.70-0.75 | 0.61-0.73 | 0.34-0.45 | 0.36-0.45 | 0.03-0.10 |
| $BC$ | 0.74-0.80 | 0.68-0.78 | 0.68-0.73 | 0.43-0.50 | 0.29-0.39 | 0.00-0.00 |
| $s$ | 0.91-0.94 | 0.86-0.89 | 0.82-0.89 | 0.77-0.83 | 0.71-0.77 | 0.17-0.28 |
| $s'$ | 0.88-0.91 | 0.86-0.91 | 0.86-0.90 | 0.76-0.79 | 0.67-0.75 | 0.21-0.32 |
| $\Delta s$ | 0.87-0.91 | 0.86-0.89 | 0.83-0.89 | 0.75-0.79 | 0.71-0.78 | 0.20-0.32 |
| $k$ | 0.70-0.74 | 0.70-0.75 | 0.57-0.69 | 0.24-0.38 | 0.23-0.36 | 0.01-0.07 |
| $k_{bu}$ | 0.90-0.92 | 0.85-0.87 | 0.77-0.83 | 0.70-0.76 | 0.68-0.77 | 0.15-0.23 |
| $k_{td}$ | 0.81-0.84 | 0.80-0.83 | 0.73-0.83 | 0.67-0.71 | 0.48-0.57 | 0.18-0.30 |
| $k_{dir}$ | 0.65-0.70 | 0.63-0.68 | 0.54-0.64 | 0.17-0.33 | 0.13-0.28 | 0.03-0.07 |
| $k_{indir}$ | 0.72-0.76 | 0.70-0.75 | 0.60-0.71 | 0.39-0.47 | 0.32-0.45 | 0.01-0.07 |
| $TI^1$ | 0.80-0.83 | 0.77-0.81 | 0.61-0.72 | 0.46-0.53 | 0.41-0.54 | 0.12-0.17 |
| $TI^3$ | 0.85-0.88 | 0.79-0.83 | 0.67-0.78 | 0.52-0.62 | 0.52-0.57 | 0.15-0.23 |
| $TI^5$ | 0.87-0.89 | 0.80-0.83 | 0.70-0.79 | 0.52-0.63 | 0.52-0.58 | 0.15-0.23 |
| $WI^1$ | 0.80-0.84 | 0.77-0.81 | 0.61-0.76 | 0.45-0.54 | 0.40-0.53 | 0.11-0.19 |
| $WI^3$ | 0.85-0.88 | 0.79-0.83 | 0.69-0.78 | 0.51-0.61 | 0.50-0.57 | 0.14-0.23 |
| $WI^5$ | 0.87-0.90 | 0.79-0.83 | 0.67-0.81 | 0.52-0.64 | 0.50-0.58 | 0.15-0.23 |
| $STO^1$ | 0.89-0.92 | 0.77-0.81 | 0.69-0.79 | 0.57-0.69 | 0.56-0.62 | 0.05-0.10 |
| $STO^3$ | 0.89-0.91 | 0.77-0.82 | 0.65-0.79 | 0.55-0.65 | 0.56-0.61 | 0.04-0.08 |
| $STO^5$ | 0.87-0.90 | 0.75-0.80 | 0.67-0.78 | 0.56-0.62 | 0.55-0.64 | 0.05-0.09 |
| $wSTO^1$ | 0.85-0.89 | 0.79-0.83 | 0.67-0.78 | 0.56-0.62 | 0.53-0.63 | 0.05-0.13 |
| $wSTO^3$ | 0.84-0.87 | 0.79-0.84 | 0.64-0.77 | 0.55-0.62 | 0.49-0.60 | 0.08-0.14 |
| $wSTO^5$ | 0.83-0.86 | 0.79-0.85 | 0.66-0.78 | 0.56-0.63 | 0.45-0.57 | 0.07-0.14 |

Table S1. Confidence intervals of the mean Kendalls in Table 1 of the paper. Green = hierarchical clustering with Jaccard index, red = hierarchical clustering with REGE index, grey = density-based modules, yellow = prey-based modules, blue = predator-based modules, purple = groups produced by the group model. C1 = Best clustering, C2 = second best clustering, C3 = third best clustering, C4 = fourth best clustering, C5 = fifth best clustering, C6 = sixth best clustering.

| Index | C1 | C2 | C3 | C4 | C5 | C6 |
|---|---|---|---|---|---|---|
| $DC$ | 50% | 50% | 50% | 50% | 25% | 25% |
| $wDC$ | 50% mean | 50% mean | 25% sum | NMAX mean | NMAX mean | NMAX sum |
| $CC$ | 50% | 50% | 50% | 50% | 25% | 25% |
| $BC$ | 50% | 50% | 50% | 25% | 25% | 25% |
| $s$ | NMAX | NMAX | 25% | NMAX | 25% | NMAX |
| $s'$ | 25% | 25% | NMAX | 25% | NMAX | 25% |
| $\Delta s$ | 25% | NMAX | 25% | 25% | NMAX | NMAX |
| $k$ | 50% | 50% | 50% | 25% | 25% | NMAX |
| $k_{bu}$ | 25% | 50% | 25% | 25% | 25% | NMAX |
| $k_{td}$ | 25% | 50% | 50% | 25% | 25% | 25% |
| $k_{dir}$ | 50% | 50% | 50% | 50% | 50% | 25% |
| $k_{indir}$ | 50% | 50% | 50% | 25% | 25% | NMAX |
| $TI^1$ | 50% | 50% | 50% | NMAX | 25% | NMAX |
| $TI^3$ | 50% | NMAX | 50% | NMAX | NMAX | NMAX |
| $TI^5$ | 25% | 25% | 25% | NMAX | NMAX | NMAX |
| $WI^1$ | 50% mean | 50% mean | 50% mean | NMAX max | 25% mean | NMAX sum |
| $WI^3$ | 50% mean | NMAX mean | 50% mean | NMAX mean | NMAX mean | NMAX mean |
| $WI^5$ | 25% mean | 25% mean | 25% mean | NMAX mean | NMAX mean | NMAX mean |
| $STO^1$ | 25% | 50% | 50% | 25% | 25% | 25% |
| $STO^3$ | 25% | 50% | 50% | 25% | 25% | 25% |
| $STO^5$ | 25% | 50% | 50% | 25% | 25% | 25% |
| $wSTO^1$ | 25% mean | 50% max | 50% mean | 25% min | 25% mean | 25% min |
| $wSTO^3$ | 25% mean | 50% mean | 50% max | 25% mean | 25% mean | 25% max |
| $wSTO^5$ | 25% mean | 50% mean | 50% mean | 25% mean | 25% mean | NMAX sum |

Table S2. Combination of linkage method and interaction strength method that gave us the highest Kendall's correlation coefficient in Table 1 of the paper. Most of the cells do not containi the interaction strength method because they are computed upon a binary network. Green = hierarchical clustering with Jaccard index, red = hierarchical clustering with REGE index, grey = density-based modules, yellow = prey-based modules, blue = predator-based modules, purple = groups produced by the group model. C1 = Best clustering, C2 = second best clustering, C3 = third best clustering, C4 = fourth best clustering, C5 = fifth best clustering, C6 = sixth best clustering.
