## Supplementary material for "Food web aggregation: effects on key positions": Description of Algorithms

### Jaccard index algorithm

1. **Compute similarity.**
   Compute the Jaccard similarity between the nodes by using the following equation (Yodzis & Winemiller, 1999)

   $$J_{ij} = \frac{a}{a+b+c} \quad (1)$$

   where $J_{ij}$ is the Jaccard similarity between node $i$ and $j$, a is the number of preys and predators that $i$ and $j$ have in common, $b$ is the number of preys and predators of $i$ but not of $j$ and $c$ is the number of preys and predators of $j$ but not of $i$.
2. **Build the dendrogram.**
   Find the two most similar elements and cluster them together. Elements are intended as nodes or clusters. Of course, during the first time we run this step all the elements are nodes. Repeat until you are left with only one item, which is the final dendrogram. During this process, the similarity between two clusters can be calculated in different ways, called linkage criteria. The ones that we used were
   - The similarity between the least similar nodes, one in each cluster, known as single linkage (Frigui, 2008)
   - The similarity between the most similar nodes, one in each cluster, known as complete linkage (Frigui, 2008)
   - The mean similarity between the nodes inside the first item and the second item, known as the weighted average distance (WPGMA). See the following equation

     $$d_{(i \cup j),k} = \frac{d_{i,k}+d_{j,k}}{2} \quad (2)$$

     where $d_{(i \cup j),k}$ is the distance between $i \cup j$ (cluster that includes $i$ and $j$) and $k$.
   - The mean similarity between the nodes inside the first item and the second item but taking into consideration the average distance between the items inside the first cluster, known as the unweighted average distance (UPGMA). See the following equation

     $$d_{(i \cup j),k} = \frac{|i|d_{i,k}+|j|d_{j,k}}{|i|+|j|} \quad (3)$$

     where $|i|$ and $|j|$ are the mean distances between the elements inside $i$ and $j$, respectively.
3. **Select the dendrogram**

   After having produced a dendrogram for every linkage criterion, select the dendrogram with the highest cophenetic correlation (Sokal & Rohlf, 1962). This allows selecting the linkage criterion that produces the dendrogram that preserves the most faithfully the pairwise similarity between different elements.

4. **Cut the dendrogram.**
   Cut the dendrogram according to the maximum inconsistency of the branches, set at 0.01.

### REGE index algorithm

1. **Compute similarity.**
   Compute the similarity between nodes by using REGE, calculated by the homonym algorithm. This was originally developed in the unpublished work by White (1980, 1982, 1984) and firstly described in the literature by Borgatti & Everett (1993). It is available to be used in the software UCINET VI (Borgatti, 2002). The REGE algorithm is the following one (Jordán, Endrédi, Liu, & D'Alelio, 2018):
   a. Set the maximum number of iterations. We set 3 iterations. Each iteration produces a matrix $R_{(t)}$ where $t$ is the number of the iteration and every element $r_{(t)ij}$ is the regular equivalence between $i$ and $j$ at iteration $t$. The regular equivalence between nodes at iteration $t = 0$ is always 1.
   b. Starting from t=1, update the elements of the matrix following these sub-steps:
      i. For every predator $k$ of species $i$, find the most similar predator $m$ of species $j$ according to $R_{(t)}$. Now, set $X_{i,k,j} = R_{(t)km}$.
      ii. For every predator $m$ of species $j$, find the most similar predator $k$ of species $i$ according to $R_{(t)}$. Now, set $X_{j,m,i} = R_{(t)mk}$.
      iii. For every prey $h$ of species $i$, find the most similar prey $n$ of species $j$ according to $R_{(t)}$. Now, set $Y_{i,h,j} = R_{(t)hn}$.
      iv. For every prey $n$ of species $j$, find the most similar prey $h$ of species $i$ according to $R_{(t)}$. Now, set $Y_{j,n,i} = R_{(t)nh}$.
      v. Update the matrix R through the following equation

$$R_{(t)ij} = \frac{\sum_{k=1} X_{i,k,j} + \sum_{m=1} X_{j,m,i} + \sum_{h=1} Y_{i,h,j} + \sum_{n=1} Y_{j,n,i}}{MAX(\sum_{k=1} X_{i,k,j} + \sum_{m=1} X_{j,m,i} + \sum_{h=1} Y_{i,h,j} + \sum_{n=1} Y_{j,n,i})} \quad (4)$$

   c. Increase $t = t + 1$ and repeat step b until you reach the maximum number of iterations. The matrix of the maximum number of iterations contains the regular equivalence between nodes.
2. **Build the dendrogram.**
   The same as in the hierarchical clustering of nodes according to their Jaccard similarity index During our analysis, we used the function linkage of MATLAB, which does not include the possibility of using a similarity matrix, so we converted the similarity matrices into dissimilarity ones. This was done by following what was written in Von Luxburg (2004). Namely, if the similarity function is normalised (takes values between 0 and 1) and always positive, then $d = 1 - s$ – where $d$ is the dissimilarity measure and $s$ is the similarity measure.
3. **Select the dendrogram.**
   The same as in the hierarchical clustering of nodes according to their Jaccard similarity index.
4. **Cut the dendrogram.**
   The same as in the hierarchical clustering of nodes according to their Jaccard similarity index.

### Topological importance algorithm

1. Compute the one step matrix.
   In the one-step matrix, if the energy flows from a prey to the predator, then the effect of the prey on the predator is the reciprocal of the indegree of the predator

$$a_{1,ij} = \frac{A_{ij}}{D_j} \quad (5)$$

2. Compute the n-step matrices.
   In the higher steps' matrices, a node influences another node at a higher trophic level by summing the effects of every path that connects the two nodes. The effect of every path is the multiplication of the inverse of the outdegree of every node along the path. For a visual explanation see Figure 2. It can be calculated as follows:

$$A(n) = A_{(1)}^{n} \quad (6)$$

3. Calculate topological importance.
   The topological importance of a node $i$ ($TI_i$) can be calculated through the following formula

$$TI_i = \frac{\sum_{m=1}^{N} \sum_{j=1}^{n} a_{m,ij}}{N} \quad (7)$$

   where N is the total number of steps considered, m is the step number, n is the total number of nodes and $a_{m,ji}$ is the effect of species i on species j at m number of steps.

Topological importance can be also used for weighted networks - giving us weighted topological importance ($WI$) – if instead of using the degree ($D$), we use the weighted degree ($WD$) (Scotti, Podani, & Jordán, 2007)

$$a_{(1)ij} = \frac{A_{ij}}{WD_j} \quad (8)$$

where $Aij$ is the element of the adjacency matrix of the weighted directed network.

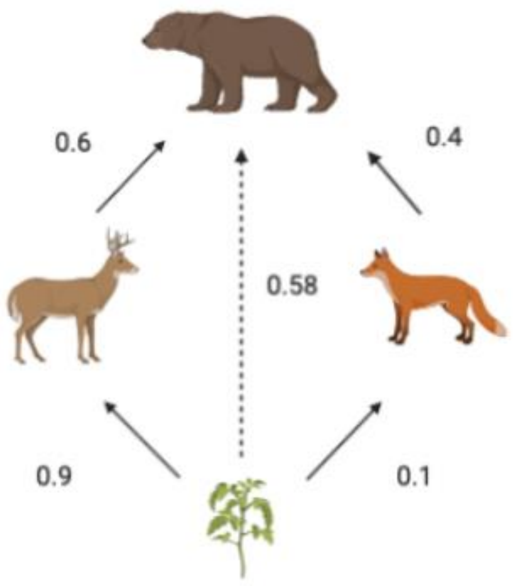


Figure A1: Topological importance (TI) of a species on another. The effects of a prey on its predator are the sum of its effects along different paths. In this case, this plant species reaches the bear through two different paths. The first one is through the deer and the second is through the fox. The first path has an effect on the bear that is 0.9*0.6=0.54, the second one has an effect on the bear that is 0.1*0.4=0.04. Summing the effects through all the paths connecting the plant with the bear, we get the effect of the plant on the bear: 0.54+0.04=0.58. Figure created with BioRender.com.

### Trophic field overlap algorithm

The trophic field overlap (TO) represents how redundant the strong interactions of a node are. It was firstly introduced by Jordán, Liu, & Mike (2009). It is the number of times that it and another node interact strongly with the same predator. The algorithm for its computation is the following one (Jordán et al., 2018):

1. Compute the one-step matrix as in topological importance.
2. Compute the n-step matrix as in topological importance.
3. Compute the average effect matrix.
   The average effect matrix ($E(n)$) represents the effect of each node on the other nodes averaged by the number of steps

$$E_n = \frac{1}{n}\sum_{i=1}^{n} A_{(i)} \quad (9)$$

4. Compute the interactor matrix.
   Compute the so-called interactor matrix ($M_T$), whose values tell us whether the interaction between two nodes is weak (W) or strong (S). To do this, we need to define a threshold over which a certain interaction is strong.
5. Compute the topological overlap matrix.
   Compute a matrix with how many times two species interact strongly with the same predator, called the topological overlap matrix.

6. Compute the trophic field overlap ($TO$).
   The trophic field overlap ($TO$) of a node is calculated by summing the elements of the rows of the topological overlap matrix.

Trophic field overlap can be also used for weighted networks – giving us weighted trophic field overlap ($WO$) – if instead of using the degree ($D$) we use the weighted degree, (e.g., Xiao et al. (2019))

$$a_{1,ij} = \frac{A_{ij}}{D_j} \quad (11)$$

### Algorithm for the deletion of simple cycles

1. Repeat the two following steps until no further cycle is left.
2. Find simple cycles.
3. Consider the shortest cycles. For each cycle, delete the weakest link inside the cycle.
