## Supplementary material for "Food web aggregation: effects on key positions": Caption Figure S1-S20

Figure S1
Upper confidence intervals of the mean Kendall's rank correlation coefficients of Figure 1. Each value is the difference between the mean and the upper confidence interval (value = upper confidence interval - mean).

Figure S2
Lower confidence intervals of the mean Kendall's rank correlation coefficients of Figure 1. Each value is the difference between the mean and the lower confidence interval (value = mean - lower confidence interval).

Figure S3 - S8
Sections of Figure 1 of the paper, which allow to read the single values in the heat map. Figure S3 = hierarchical clustering using Jaccard index, Figure S4 = hierarchical clustering using REGE index, Figure S5 = maximisation of density-based modularity, Figure S6 = maximisation of prey-based modularity, Figure S7 = maximisation of predator-based modularity, Figure S8 = group detection through the group model.

Figure S9 - S14
Sections of Figure S1, which allow to read the single values in the heat map. Figure S9 = hierarchical clustering using Jaccard index, Figure S10 = hierarchical clustering using REGE index, Figure S11 = maximisation of density-based modularity, Figure S12 = maximisation of prey-based modularity, Figure S13 = maximisation of predator-based modularity, Figure S14 = group detection through the group model.

Figure S15 - S20
Sections of Figure S2, which allow to read the single values in the heat map. Figure S15 = hierarchical clustering using Jaccard index, Figure S16 = hierarchical clustering using REGE index, Figure S17 = maximisation of density-based modularity, Figure S18 = maximisation of prey-based modularity, Figure S19 = maximisation of predator-based modularity, Figure S20 = group detection through the group model.
